# Molecular mechanisms underlying Warburgia salutaris effects on oxidative stress and apoptotic parameters in Human Hepatoma Cells

**DOI:** 10.1101/2022.03.05.483129

**Authors:** Lebogang N. Maruma, Anou M. Somboro, Daniel G. Amoako, Hezekiel M. Khumalo, Rene B. Khan

## Abstract

This study aims to determine the molecular effects of *Warburgia salutaris* extract in HepG2 cells and elucidate the possible mechanisms. The MTT assay was employed to determine cell viability and the half maximal inhibitory concentration (IC_50_) of *Warburgia salutaris-treated* in HepG2 cells (0-5mg/ml). Extracellular lactate dehydrogenase and ATP were also quantified as a measure of cell viability. The production of reactive oxygen species (ROS) was assessed by quantifying lipid peroxidation and oxidative DNA damage, and reactive nitrogen species (RNS) in treated HepG2 cells. The cells response to free radicals was assessed by measuring GSH. Stress response antioxidant and apoptotic markers were detected using western blotting and /or qPCR. Cell death parameters assayed included annexin V, caspase activity and necrosis. Single-cell gel electrophoresis (SCGE) was used to visualise DNA damage in the HepG2 cells and confirmed with DNA fragmentation assay. The Hoechst assay allowed the visualisation of the nucleus to assess cell growth and apoptosis. Decreased cell viability was associated with a decreased level of ATP. The presence of oxidative stress was suggested by increased HSP70 and Nrf2 protein expression and confirmed by increase ROS, RNS, GPx and catalase; and a corresponding decrease of SOD2 and glutathione. Caspase 8 showed no significant difference between treatment concentrations, caspase 9 was decreased and caspase 3/7 increased. A reduction in p53 correlated with chromatin changes, increase in comet lengths and DNA fragmentation. NFκB protein was significantly decreased at the IC_50_, along with decreased cMyc protein expression. Our findings shows that *Warburgia salutaris* promotes apoptosis by inducing oxidative stress in HepG2 cells and may be a potential anti-cancer agent that would serve as an alternative to conventional therapeutic agents.

## 1. Introduction

Cancer is a non-communicable disease, and the second cause of death in most countries following cardiovascular diseases ^1^. In 2018 alone, approximately 18. 1 million new cancers cases and a death toll of 9.6 million was reported ^2^. Liver cancer is the seventh most common cancer worldwide in terms of incidence, and fourth leading cause of all cancer-related mortality in both sexes^2,3^. In the same year, an estimate of 841 000 new cases and a global death of 782 000 was associated with liver cancer. It is more common in males than females; and males have 4-8-fold more likely to develop liver cancer than females, ranking liver cancer as the second leading cause of cancer-related mortality in men. Compare to males, females have higher chances of survival ^2,4,5^

Hepatocellular carcinoma (HCC) and intrahepatic cholangiocarcinoma (ICC) are two main histologic types of liver cancer. HCC is the dominant subtype representing 75-80% of all types of liver cancer. Hepatitis B, hepatitis C, and diary aflatoxin, most notably aflatoxin B_1_ (AFB_1_) are the major risk factors associated with 80% of HCC. Smoking, excessive alcoholic consumption, and non-alcoholic fatty liver disease related to diabetes and obesity are the other minor risk factors. Majority of the new cases are found in resource constrained regions such as Sub-Saharan Africa and Eastern Asia ^6–8^. HCC may be treated using chemotherapy, radiotherapy and/or surgical resection ^9^. However, the prognosis for liver cancer remains poor, and 92% of diagnosed patients die within 1 year of the onset of symptoms. Outcomes are poor, with an estimated 5-year net survival rate of 19%^10^. In South Africa, approximately 46 000 new liver cancer cases are diagnosed annually, with an increase in incidence of 41.2 per 100 000 persons per year ^9^. It is clear, based on poor prognosis and low survival rates, that existing treatment options are ineffective and need expanded research to find new effective HCC therapies.

For millennia, plant extracts have been exploited for the treatment of a wide variety of illnesses. Current research trends present traditional herbal medicine as a significant biomedical resource that may contribute significantly to the treatment of cancer. Traditional herbal medicines are already used extensively by 80% of the rural African population to treat various diseases, including cancer^11^. For example, *Sutherlandia frutescens* (cancer bush) is utilised by South African traditional healers to treat cancer as it possesses anti-proliferative effects in lung cancer cells ^12^. Leshwedi demonstrated the potential of methanolic extracts of *Warburgia salutaris* to protect peripheral blood mononuclear cells against the carcinogenic effects of crystalline silica ^13^. Further support for the use of *Warburgia salutaris* as a therapeutic agent against cancer was provided by Soyingbe *et al*. (2018), where an anti-proliferative effect in MCF-7 cells was attributed to the acetone fraction of *Warburgia salutaris* ^14^.

*Warburgia salutaris* is a herbal remedy used by traditional healers for treating medical sicknesses such as coughs, colds, pleural infections, oral thrush, and cystitis ^15^. It is also exploited for its antioxidant properties that may be conferred by the flavonoids and flavonols present ^16^. Thus, *Warburgia salutaris* may possess both antioxidant and anti-proliferative activity against cancerous cells. There is currently no documented evidence of the effects of *Warburgia salutaris* on HCC cells, and its mechanisms have not been fully elucidated. Therefore, this study aims to investigate the molecular effects of *Warburgia salutaris* in HepG2 cells.

## 2. Materials and methods

### 2.1. Materials

The human hepatocellular carcinoma (HepG2) cell line was purchased from Highveld biologicals (Johannesburg, South Africa). *Warburgia salutaris* capsules were obtained from a local pharmacy (Health box, shop number L13, West Wood Mall, Westville) in Durban. All the cell culture reagents were procured from Whitehead Scientific (Johannesburg, South Africa). Phosphate buffered saline (PBS), MTT salt, malondialdehyde (MDA), thiobarbituric acid (TBA) and β-actin were purchased from Sigma Aldrich (St Louis, MO, USA). Foetal calf serum, bovine serum albumin (BSA), gel red and all primers were purchased from Inqaba Biotech (Johannesburg, South Africa). Western blotting and PCR equipment and reagents were obtained from Bio-Rad (Hercules, CA, USA). Cell Signalling Technology antibodies and Promega luminometry kits were procured from Anatech (Johannesburg, South Africa). All other reagents were purchased from Merck (Darmstadt, Germany), except where otherwise stated.

### 2.2. Preparation of *Warburgia salutaris* extract

*Warburgia salutaris* (30 capsules) were extracted by adding 100ml of distilled water and stirring for 2 hours before transfer into two 50ml tubes. The tubes were centrifuged (Eppendorf, Germany) for 5 minutes at 2000×g. The extract supernatants were poured into two freeze dryer bottles and frozen at −20°C for 24 hours. After 24 hours, the extract was freeze dried (Edwards Monor Royal, Czech) completely and percentage yield was calculated. The extract was stored at 4°C until used. A crude *Warburgia salutaris* (10mg/ml) stock solution was prepared fresh as required. Serial dilutions for the MTT assay (0-5mg/ml), as well as the inhibitory concentrations were prepared from the stock solution.

### 2.3. Cell culture

HepG2 cells were cultured in complete culture medium (CCM) comprising of Eagle’s minimum essential medium (EMEM) supplemented with 10% foetal calf serum, 1% penicillin-streptomycin-fungizone and 1% L-glutamine at 37°C in 5% CO_2_ incubator. Cells were washed every two days with 0.1M phosphate buffered saline (PBS, pH 7.4) and replenished with 5 ml of CCM. When cells reached 90% confluence, the cells were washed in 0.1ml PBS, detached by incubation in 1ml trypsin at 37°C for 5 minutes. The number of viable cells were counted using the Trypan blue cell exclusion method.

### 2.4. MTT assay

The MTT assay was used to determine the cytotoxicity of *Warburgia salutaris* extract on HepG2 cells. HepG2 cells (15000 cells/well) were cultured in a 96-well microtitre plate (200μL) and treated with *Warburgia salutaris* crude water extract at concentrations ranging from 0-5mg/ml for 24 hours. Subsequently, the treatment medium was discarded, the cells were added with 20μl MTT salt solution (5mg/ml in 0.1 ml PBS) and 100μl CCM per well. After incubation for 4 hours at 37°C, the MTT salt solution was discarded and 100μl of dimethyl sulphoxide (DMSO) was added into each well and incubated for 1 hour at 37°C. The optical density of the solubilised formazan product was detected using a spectrophotometer (Bio-tek μQuant, USA) at 570nm/690 nm. The results were reported as

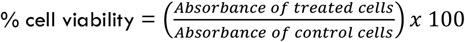

A graph was generated by plotting cell viability and the concentration-dependent response using GraphPad Prism v5.0 software (GraphPad software Inc, La Jolla, California). A maximum inhibitory concentration of 50% (IC_50_: 4.8mg/ml) was obtained for use in all subsequent assays. In addition, cells were also treated with half the IC_50_ concentration (IC_25_). IC_50_ was chosen as a half-maximum inhibitory concentration to measure the potency of *Warburgia salutaris* in HepG2 and IC_25_ was used to measure the potency of *Warburgia salutaris* at lower doses in HepG2.

### 2.5. Thiobarbituric acid reactive substances (TBARS) assay

HepG2 cells were cultured in tissue culture flasks (400 000 cells) and treated with 2.4mg/ml (IC_25_), 4.8mg/ml (IC_50_) and CCM as a control for 24 hours. Following treatment, an amount of 200μL of supernatant per control/treated were resuspended in 0.2% phosphoric acid (H_3_PO_4_), homogenised by passing 25 times through a 25 gauge needle and the homogenate transferred to a glass tube. Both negative (200μl CCM) and positive (199μl CCM + 1μl MDA) control was also prepared. Subsequently, 200μl of 2% H_3_PO_4_ and 200μl of 7% H_3_PO_4_ was added to each of the glass test tubes. Thereafter, each sample received 400μl of TBA/Butylated hydroxytoluene (BHT) solution, except the blank to which 400μl of 3mM HCl was added. Each tube was vortexed for few minutes and the pH of each sample acidified by adding 200μl of 1 M HCl. The samples were first boiled to 100°C in a water bath for 15 minutes, then cooled to room temperature. Butanol (1500μl) was added to each tube and vortexed for 30 seconds. The upper 1000 μl of the butanol phase was extracted and transferred into 1500μl micro-centrifuge tubes and centrifuged at 2500 x g speed at 24°C temperature for 6 min. Each sample was added into triplicate wells of a 96-well microtitre plate (200μl) and absorbance was measured using a Bio-Tek μQuant spectrophotometer (USA) at 532nm, with a reference wavelength of 600nm. Average MDA concentration was calculated using the following equation:

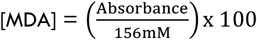

### 2.6. Nitrates assay

Treated HepG2 cells were then trypsinised, counted and resuspended in PBS. Nitrate standards were prepared (0μM–200μM) and each standard (50μl) and sample supernatant (50μl) were added into a 96 well microtiter plate in triplicates. Next, the following reagents were added in quick succession into each well; 50μl VCl_3_, 25μl SULF and 50μl NEDD. The mixture was incubated for 45 minutes at 37°C and absorbance was measured using a Bio-Tek μQuant spectrophotometer (USA) at 540nm, with a reference wavelength of 690nm. The standard curve produced was used to determine RNS in the samples.

### 2.7. Luminescent Assays

CellTiter-Glo^®^ luminescent cell viability assay (Promega, Madison, Wisconsin) was used to quantify adenosine triphosphate (ATP). The treated HepG2 cells (20 000 cells in 200μl CCM) were pipetted in duplicates into an opaque-walled multiwell plate. Thereafter, the treatment media was removed, and the cells were washed once with PBS. The wells were replenished with 50μl PBS and 25μl CellTiter-Glo^®^ reagent (Cat. #G7570, prepared as per the manufacturer’s instructions). The contents were mixed on an orbital shaker and the plate was incubated at room temperature for 30 minutes. Luminescence was recorded in relative light units (RLU) using a Modulus™ microplate luminometer (Turner Biosystems, Sunnyvale, USA).

### 2.8. Measurement of Caspase Activity

Caspase-Glo^®^ assay was used to measure the activity of caspase-8, −9 and −3/7. In brief, 50μl of treated HepG2 cells (20 000 cells, 200μl CCM) were seeded into wells of an opaque multiwell plate. Caspase 8-Glo^®^ reagent (Cat. #G8200) was prepared by following manufacturer’s instruction and 25μl was added to each well. After a brief mix on the orbital shaker, the plate was incubated at room temperature for 30 minutes. The Modulus™ microplate luminometer (Turner Biosystems, Sunnyvale, USA) was used to record luminescence in RLU. This procedure was repeated for caspase 9 (Cat. #G8210) and caspase 3/7 (Cat. #G8090).

### 2.9. Measurement of apoptosis and necrosis

The RealTime-Glo^™^ Annexin V Apoptosis and Necrosis Assay was used to assess apoptosis and necrosis in treated HepG2 cells. The HepG2 cells (20 000 cells, 200μl/well) were plated into duplicate wells of a white multiwell plate. Following treatment, the cells were washed with PBS, then 50μl PBS was added to each well. The 2X detection reagent was prepared according to the manufacturer instruction and 25μl of this mixture was then added into each well. The contents were mixed briefly on an orbital shaker and the plate was incubated at room temperature for 30 minutes. Apoptosis and necrosis were quantified using the Modulus™ microplate luminometer (Turner Biosystems, Sunnyvale, USA) and measured in RLU for apoptosis and relative fluorescence units (RFU) for necrosis, respectively.

### 2.10. Glutathione (GSH)

The HepG2 cells (20 000 cells/well, 200μl CCM) seeded in duplicate in an opaque white-walled microtitre plate were treated for 24 hours. The GSH-Glo^™^ reagent (25μl) was added to the culture, along with 50μl PBS. The contents were mixed on an orbital shaker and the plate was incubated at room temperature for 30 minutes. Thereafter, the luciferase was added, and the plate was incubated for a further 15 minutes before recording the luminescent signal in RLU (Modulus™ microplate luminometer, Turner Biosystems, Sunnyvale, USA).

### 2.11. Lactate dehydrogenase (LDH) assay

The Roche LDH assay was used to measure the release of LDH from damaged cell membrane. Treatment of HepG2 cells (20 000 cells/well in 200μl CCM) in duplicate wells of microtitre plate for 24 hours was followed by the transfer of the treatment medium into a new multiwell plate. The LDH reagent (100μl) was added to each well. After 30 minutes, stop solution (50μl) was added. The absorbance was recorded using a spectrophotometer (Bio-tek μQuant, USA) at 490nm/600nm.

### 2.12. Comet assay

The comet assay was used to measure the degree of DNA fragmentation. Treated HepG2 cells were embedded in LMPA slides with frosted ends. The first layer was formed by adding 2% low melting point agarose (800 μl, 37°C) to a slide, covered with a coverslip on top and gel allowed to solidify at 4°C for 10 minutes. After removing the coverslip, the second layer containing cells (20 000 cells in 25μl PBS), gel red (1μl) and 1% LMPA was added to the first layer. The gel was covered with a coverslip and left to solidify at 4°C for 10 minutes. Finally, the coverslip was removed and the third layer (1% LMPA, 200μl; 37°C) was added and overlayed with a coverslip. Once the gel was solidified at 4°C for 10 minutes, the coverslip was removed. Subsequently the slides were covered with cold cell lysis buffer [100mM EDTA, 2.5M NaCl, 1% Triton X-100, 10% DMSO, and 10mM Tris (pH 10)] for 1 hour at 4°C. Afterwards, the lysis buffer was removed, and the slides were submerged in electrophoresis buffer (1mM Na2EDTA and 300mM NaOH, pH 13) for 20 minutes at room temperature for equilibration. Electrophoresis (25V, 35 minutes) proceeded using a Bio-Rad compact power supply. The cells were rinsed three times (5 minutes each) with 0.4M Tris, (pH 7.4) to neutralise the samples. The slides were viewed using fluorescent microscope (Olympus IX51 inverted microscope, excitation: 510-560 nm; emission 590 nm). Images of 50 comets were captured and measured (Life Science - ©Olympus Soft Imaging).

### 2.14. Hoechst assay

Hoechst 33342 (H3570; Invitrogen, Eugene, Oregon) was used to assess the nuclear morphology in HepG2 cells treated with *Warburgia salutaris.* HepG2 cells were plated in two 6-well plates (400 000 cells/well) in duplicate for each treatment. The cells were treated with 2.4 mg/ml (IC_25_), 4.8 mg/ml (IC_50_), 7.2 mg/ml (IC_75_) and CCM as a control for 24 hours. The cells were washed with PBS. The Hoechst working solution (5μl in 10ml PBS) was prepared and 500μl was added into each well. The plate was incubated for 15 minutes (37°C) and the cells were washed with PBS. Subsequently, the cells were then fixed using 10% paraformaldehyde (pH 7.4) for 5 minutes. The cells were washed again and viewed using the 350nm/461nm excitation/emission filter of the Olympus Microscope Life Science - ©Olympus Soft Imaging Solutions v5 to detect nuclear changes.

### 2.15. Western blotting assay

After treatment, total protein was extracted using CytoBuster^™^ reagent, supplemented with protease and phosphatase inhibitors according to manufacturer’s protocol. The pellet was discarded, and the retained lysate was quantified for protein concentration using the bicinchoninic acid (BCA; Sigma, St Louis, Missouri) assay. Proteins were standardized to 1mg/ml, mixed with sample buffer and denatured (100° for 5 minutes). The denatured protein was loaded onto a polyacrylamide gel (10% resolving, 4% stacking) in a tank containing electrophoresis buffer (Tris; glycine; SDS; dissolved in dH_2_O) and subjected to electrophoresis (150V, 90 minutes). The separated proteins were then transferred (Tris; glycine; SDS; dissolved in dH_2_O) onto a nitrocellulose membrane (25V, 2.5A, 30 minutes) using a Transblot^®^ Turbo Transfer system (Bio-Rad). The membrane was incubated in blocking buffer for 2 hours ((5% BSA in TTBS (NaCl; KCl; Tris; dissolved in dH_2_O)), then immunoblotted with an exact primary (Appendix 1) antibody (1:1000 in 5% BSA/TTBS) for 1 hour and placed overnight in the 4°C refrigerator. The membrane was washed 5x with TTBBS (10 minutes per wash) and then HPR conjugated secondary antibody (1:5000 in 5% BSA/TTBS Appendix 1) was added for 2 hours. After 5x 10-minute washes in TTBS, the membrane visualization was facilitated by 1:1 chemiluminescence luminol enhancer solution and peroxide solution using the Bio-Rad imaging system (Molecular Imager^®^ Chemidoc XRS and Bio-Rad imaging system).

The membranes were stripped, before the re-probing for the housekeeping protein. The membrane was washed once with distilled water, then incubated in hydrogen peroxide (10ml) for 30 minutes at 37°C. The membrane was rinsed with distilled water and then incubated with TTBS on a shaker for 5 min. Blocking proceeded by adding 5 ml 5% BSA/TTBS for 2 hours with agitation. The HRP-labelled β-actin-antibody (1:5000 in 5% BSA/TTBS) was added for 1 hour. The membrane was washed 5x with TTBBS (10 minutes per wash). The image was captured using the Chemidoc XRS and Bio-Rad framework from Molecular Imager^®^.

### 2.16. Quantitative Polymerase Chain Reaction (qPCR assay)

#### 2.16.1. RNA isolation

Triazol (500μl) and PBS (500μl) was added to each treated HepG2 cells and a scraper was used to dislodge the attached cells. Thereafter, 100μl of chloroform was put into each sample and shaken robustly for 15 seconds followed with a 2-3-minutes incubation at room temperature and centrifugation for minutes (12000×g; 4°C). The aqueous phase was transferred to a new eppendorf and 250μl was added followed by a flick to mix and storage at −80°C overnight. The thawed samples were centrifuged (20 minutes; 12000×g; 4°C), the supernatant was removed, and the pellet was washed by adding ethanol (75%; 500μl) and centrifugation (15 minutes; 12000×g; 4°C), after which the ethanol was removed, and the pellets were left to dry (1 to 1.5hours). The dry pellets were resuspended in 15μl of nuclease free water and incubated at room temperature for 2 to 3 minutes on ice. The RNA samples were then quantified using Nanodrop 2000^®^.and standardised (1000ng/μl).

#### 2.16.2. cDNA synthesis

In preparation for cDNA synthesis, 4μl of each RNA sample was added to 16μl of a prepared reaction mix (Appendix 2). The tubes were inserted into Gene Amp^®^PCR system 9700 version 3.05 and run at preset reaction conditions (Appendix 2). After the run, 80μl of nuclease free water was added, and the cDNA was stored at −80°C until needed.

#### 2.16.3. qPCR

The PCR mastermix (Sybr green, forward/reverse primer, nuclease free water) was prepared for the target gene protein (SOD2, OGG1) and housekeeping protein (GAPDH) (Appendix 3). The housekeeping and target gene protein were pipetted into a PCR plate in triplicate (11 μl), then cDNA (1.5μl) was added to the respective wells. The plate was centrifuged at 2000rpm (3 minutes at room temperature) and placed in the CFX Touch^™^ Real Time PCR Detection System (Bio-Rad; Hercules, California, USA) set for an initial denaturation (95°C, 4 minutes) and 40 cycles that included denaturation (95°C, 15 seconds), annealing (55°C, 40 seconds) and extension (72°C, 30 seconds), and a final hold step. The data obtained was represented as relative fold change in mRNA expression (2-ΔΔCT) relative to the control.

### 2.17 DNA fragmentation

Treated cells were centrifuged (1000000 cells) (Eppendorf Centrifuge 5804R, Hamburg, Germany) for 10 minutes (1491 rpm, 24°C), and the cell pellet was suspended in cell lysis solution (600μl) for 15 minutes on ice. Potassium acetate (600μl) was added to the lysed cells, vortexed and invert-mixed for 8 minutes. Half of the sample was transferred to a clean eppendorf tube, and subsequently centrifuged at 13 000rpm for 5 minutes. The addition of isopropanol (600μl) was followed by an invert mix of another 5 minutes, then centrifugation at 13 000 rpm for 5 minutes. Each pellet was resuspended in 60μl isopropanol (eppendorf), the pellets were mixed, and the final pellet has 300μl of ethanol. The ethanol was removed (supernatant) and the tubes were inverted for 15 minutes. The pellet has 40μl (10mM EDTA, pH8, 100mM Tris cl (pH 7.4) and dH20) of a DNA hydration solution when dry. Then the tubes were vortexed and placed in a water bath (set at 65°C). The DNA concentration of each sample was measured using Nanodrop (Nanodrop 2000). The tests were carried out for low-level (100ng/μl) DNA standardization. The standardized DNA was electrophoresed following the loading of specimens into 1, 8% with ethidium bromide (10mg/mL). A voltage of 120V was used and the gel was measured using the UV Tech Alliance 2.8 system (Quantity 1 measuring software). Fragmentation of DNA by Autoradiography imagery system was viewed.

### 2.18. Data Analysis

All statistical analyses were carried out using GraphPad Prism Version 5 software (GraphPad Software Inc., La Jolla, USA). Data was expressed as mean ± standard deviation. Three replicates were used, and all experiments were repeated to ensure reproducibility. One-way analysis of variance followed by a Tukey’s significant difference test was used to determine statistically significance (*p* < 0.05) differences.

## 3. Results

### 3.1. Cell Viability and Metabolic Activity

#### 3.1.1. MTT assay

The colorimetric MTT assay was used to examine *Warburgia salutaris* toxicity in HepG2 cells over 24 hours. *Warburgia salutaris* concentrations (0-5mg/ml) were used to determine a dose response activity. At the lower concentrations (< log 0.25mg/ml), the cell viability was increased relative to the control (Figure 1B). Subsequent increases in concentration resulted in a dose-dependent decrease in cell viability, with the lowest viability recorded at 5mg/ml (Figure 1B). The curve shows that 2.4mg/ml and 4.8mg/ml was sufficient to cause 25% (IC_25_) and 50% (IC_50_) cytotoxicity in HepG2 cells respectively (Figure 1A). The IC_25_ and IC_50_ concentrations were used in subsequent assays and were compared to untreated cells (CCM only).

**Figure 1:**
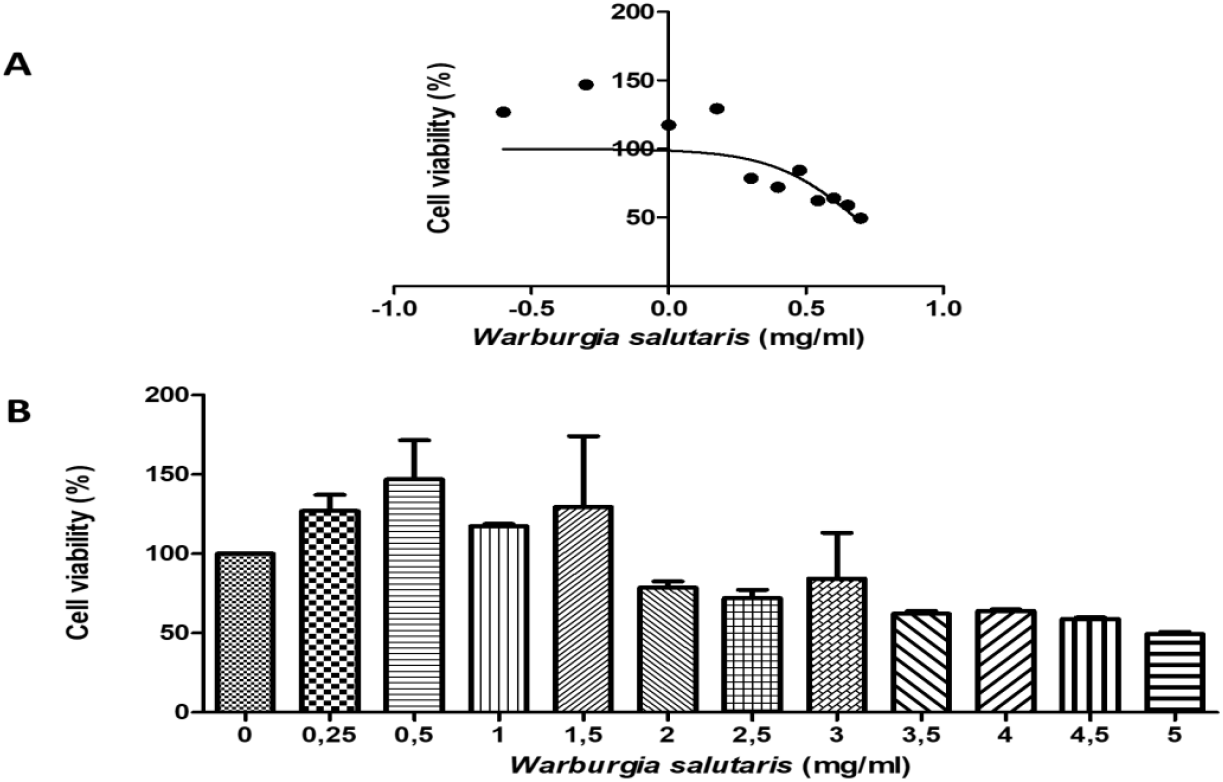
**A** The dose-response curve derived from data expressed as percentage cell viability versus log concentration of *Warburgia salutaris* was used to calculate the IC_50_ concentration. **B** Cell viability (%) of HepG2 cells treated with *Warburgia salutaris* over 24 hours. The % cell viability was increased at concentrations less than 2mg/ml, and at concentrations of >2mg/ml the % cell viability was reduced below the control (100%).

#### 3.1.2. Metabolic activity: ATP quantification assay

The ATP quantification assay was used to assess metabolic activity in the HepG2 cells treated with various concentrations (IC_25_ and IC_50_) of *Warburgia salutaris*. The ATP levels were reduced 16-fold in the IC_25_ treatment compared to the control (6850000±50000RLU). Although the ATP concentrations reduced in the IC_50_, it was not significant compared to IC_25_ (Figure 2).

**Figure 2:**
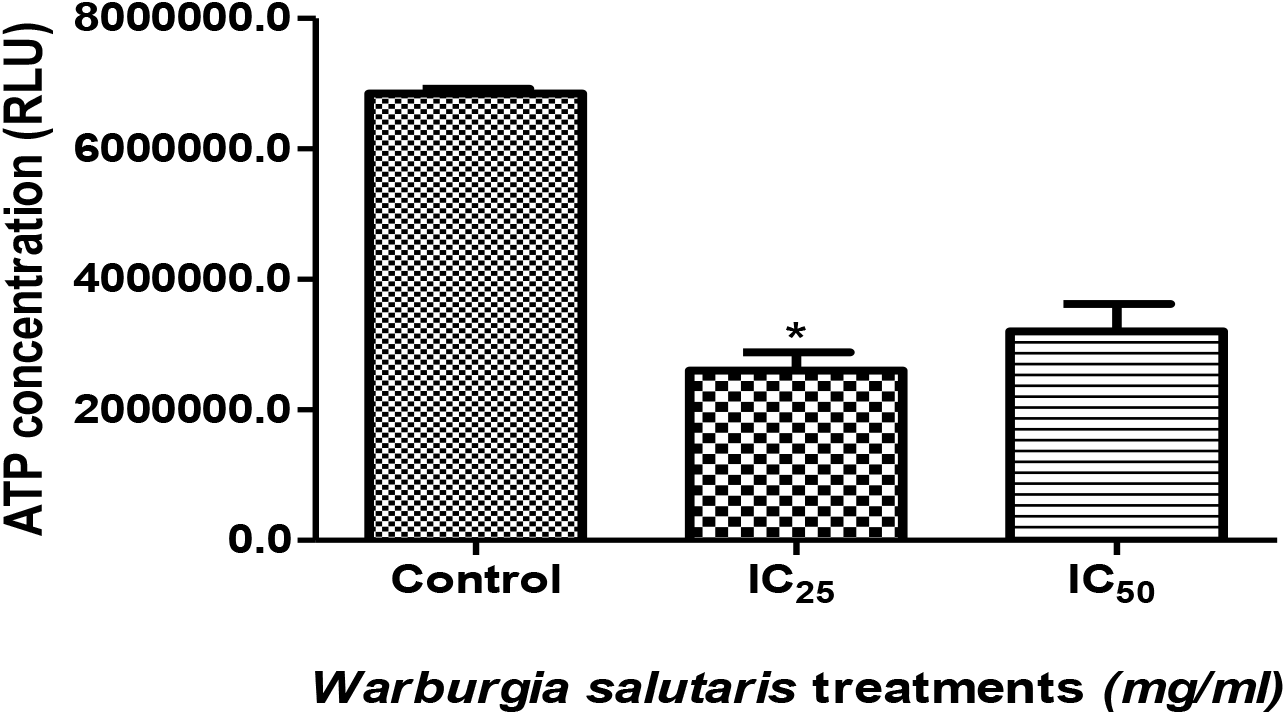
ATP concentration in HepG2 cells treated with *Warburgia salutaris* was significantly reduced for all concentrations relative to the control (\**p*=0.0309, Students *t*-test with Welch’s correction).

### 3.2. Oxidant Production/Damage

The production of ROS and RNS was quantified using the TBARS and nitrates assays, respectively. Assessment of oxidative damage to lipids and DNA was assessed.

#### 3.2.1 Nitrates assay

The NO levels produced in the HepG2 cells treated with *Warburgia salutaris* were significantly increased in the IC_25_ treatment, with no change observed in the IC_50_ (Figure 3a) compared to the control.

**Figure 3a:**
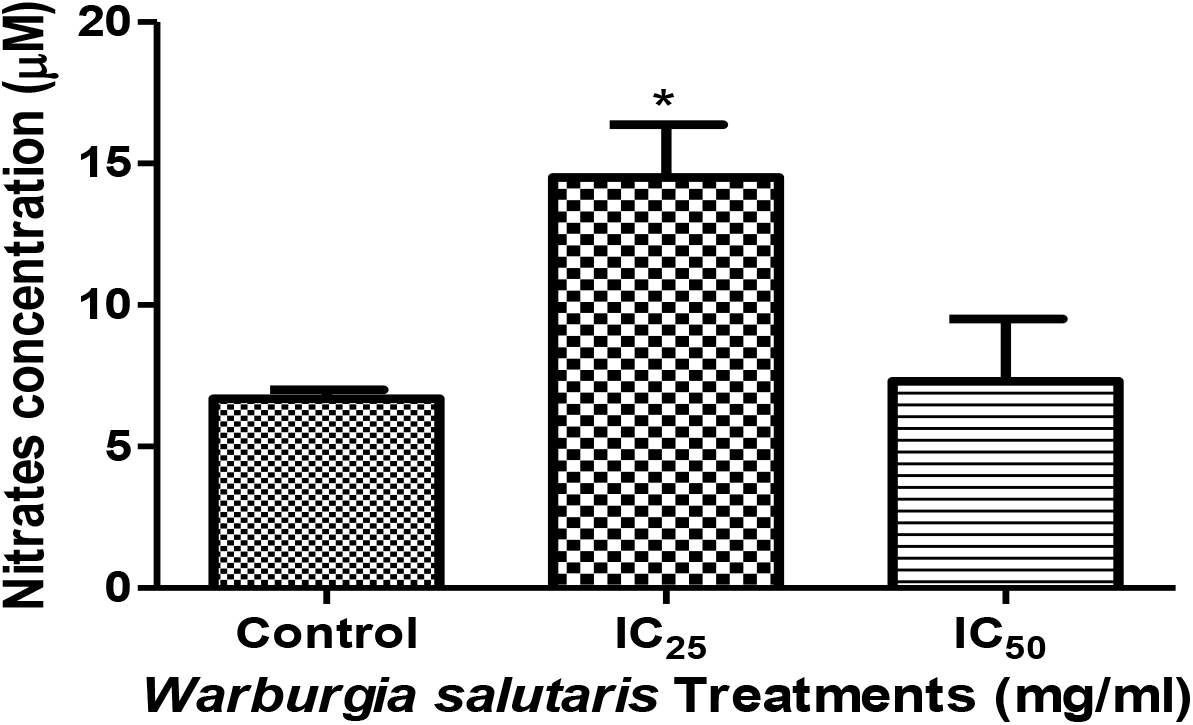
NO levels in the IC_25_-treated HepG2 cells were significantly increased (\**p*=0.0192, Students *t*-test with Welch’s correction), and the IC_50_ was similar to the control.

#### 3.2.2. TBARS assay

With regards to the TBARS assay (Figure 3b), the MDA levels determined were significantly increased in all treatment concentrations relative to the control (*p*<0.0001, ANOVA).

**Figure 3b:**
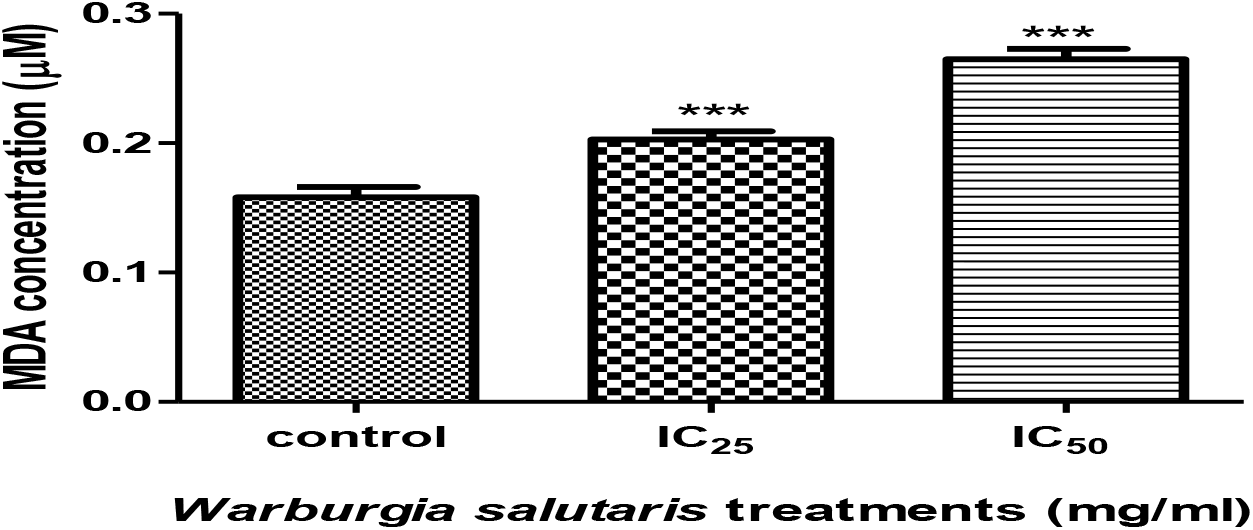
MDA concentration increased significantly in all cells treated with *Warburgia salutaris* (IC_25_, \*\*\**p*=0.0003 and IC_50_, \*\*\**p*=0.0001; Students *t*-test with Welch’s correction).

#### 3.2.3. OGG1 gene expression

The DNA repair enzyme OGG1 is a marker of oxidative DNA damage. OGG1 gene expression did not significantly decrease for the IC_25_ treatment, but there was a significant for IC_50_ treatment compared with the control (Figure 3c).

**Figure 3c:**
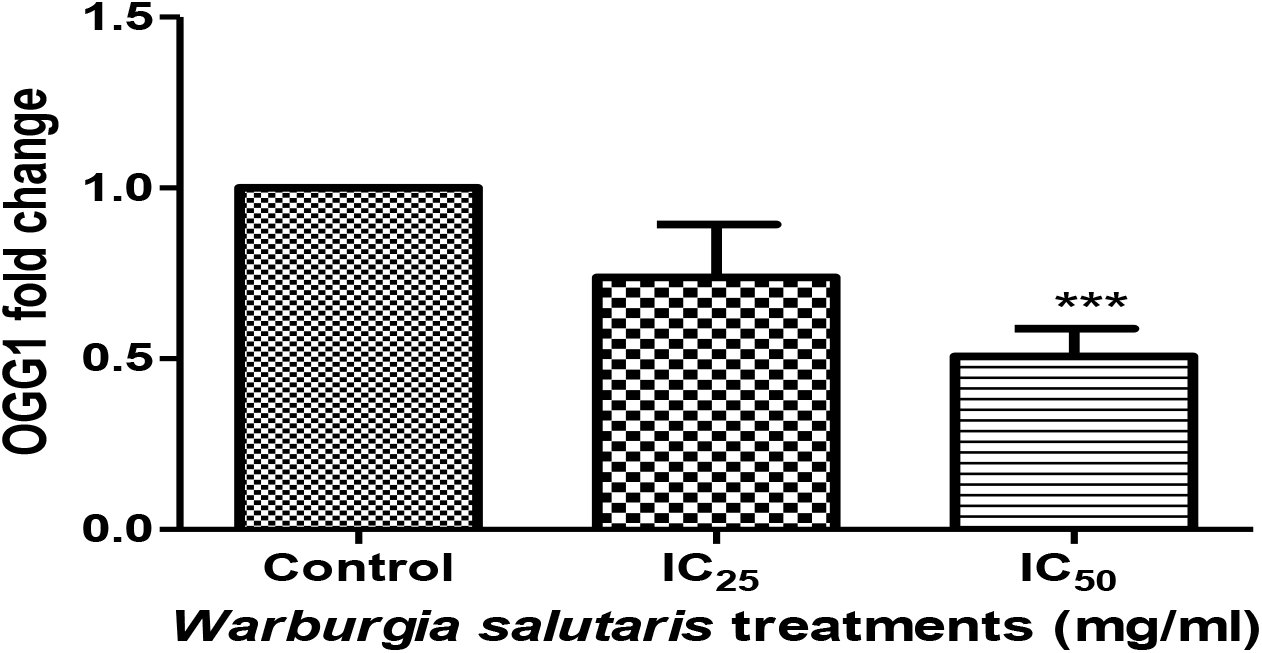
OGG1 gene expression in HepG2 cells treated with *Warburgia salutaris* shows a significant decrease at the IC_50_ (\*\*\**p*<0.05) compared with the control, Students *t*-test with Welch’s correction.

### 3.3. Antioxidant response

#### 3.3.1. GSH quantification

The GSH quantification assay was used to assess the extent of oxidative stress in the cell. GSH concentration was decreased in *Warburgia salutaris* treated cells (*p*<0.0002 ANOVA) compared to the control (Figure 3d). The results showed a decrease in GSH levels at both treatment concentrations, IC_25_ (significant) and IC_50_ (not significant).

**Figure 3d:**
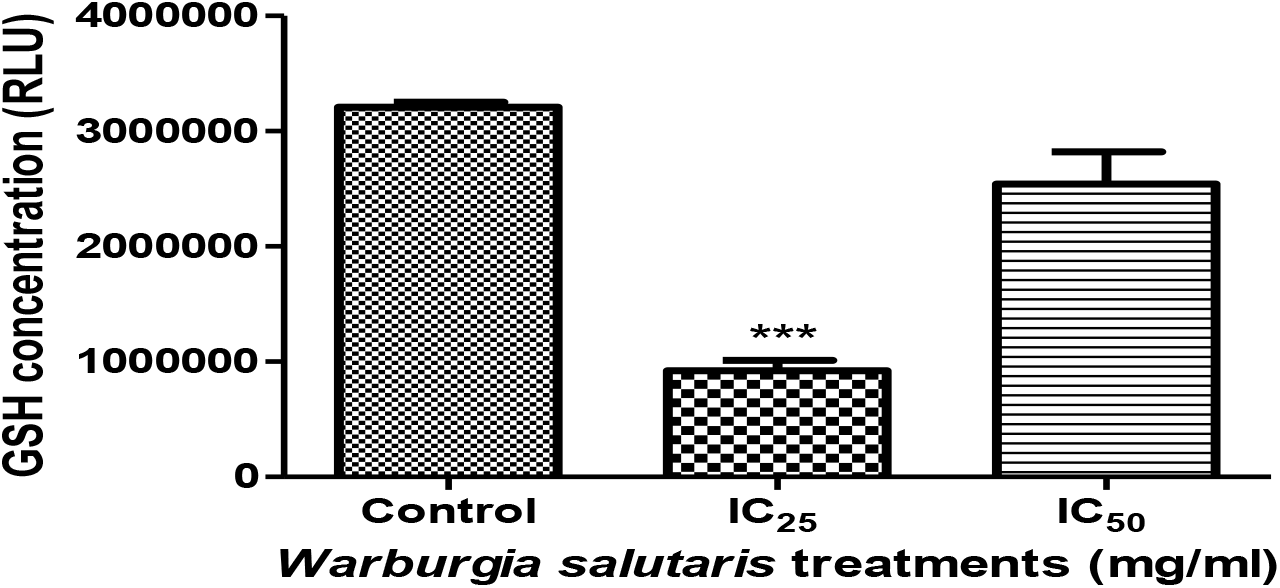
The GSH concentration in HepG2 cells treated with *Warburgia salutaris* decreased significantly at the IC_25_ treatment (\*\*\**p*=0.0007; Students *t*-test with Welch’s correction).

#### 3.3.2 Stress-related and antioxidant proteins

The effects of *Warburgia salutaris* on HepG2 stress-related and antioxidant protein expression were assessed by western blotting. All protein expression was compared to the control. The HSP70 protein expression in Figure 4A shows a significant increase at the IC_25_, while the IC_50_ showed no significant change compared to untreated cells. The Nrf2 (Figure 4B) was similarly increased at the IC_25_, while a non-significant increase was observed at the IC_50_. Oxidative stress protein markers (Figure 4C, D, E) SOD2 (Figure 4C) significantly decreased at both the IC_25_ and IC_50_. Conversely, catalase (Figure 4D) increased significantly at both concentrations, while GPx (Figure 4E) showed a non-significant increase at the IC_25_ and IC_50_ compared to the control.

**Figure 4:**
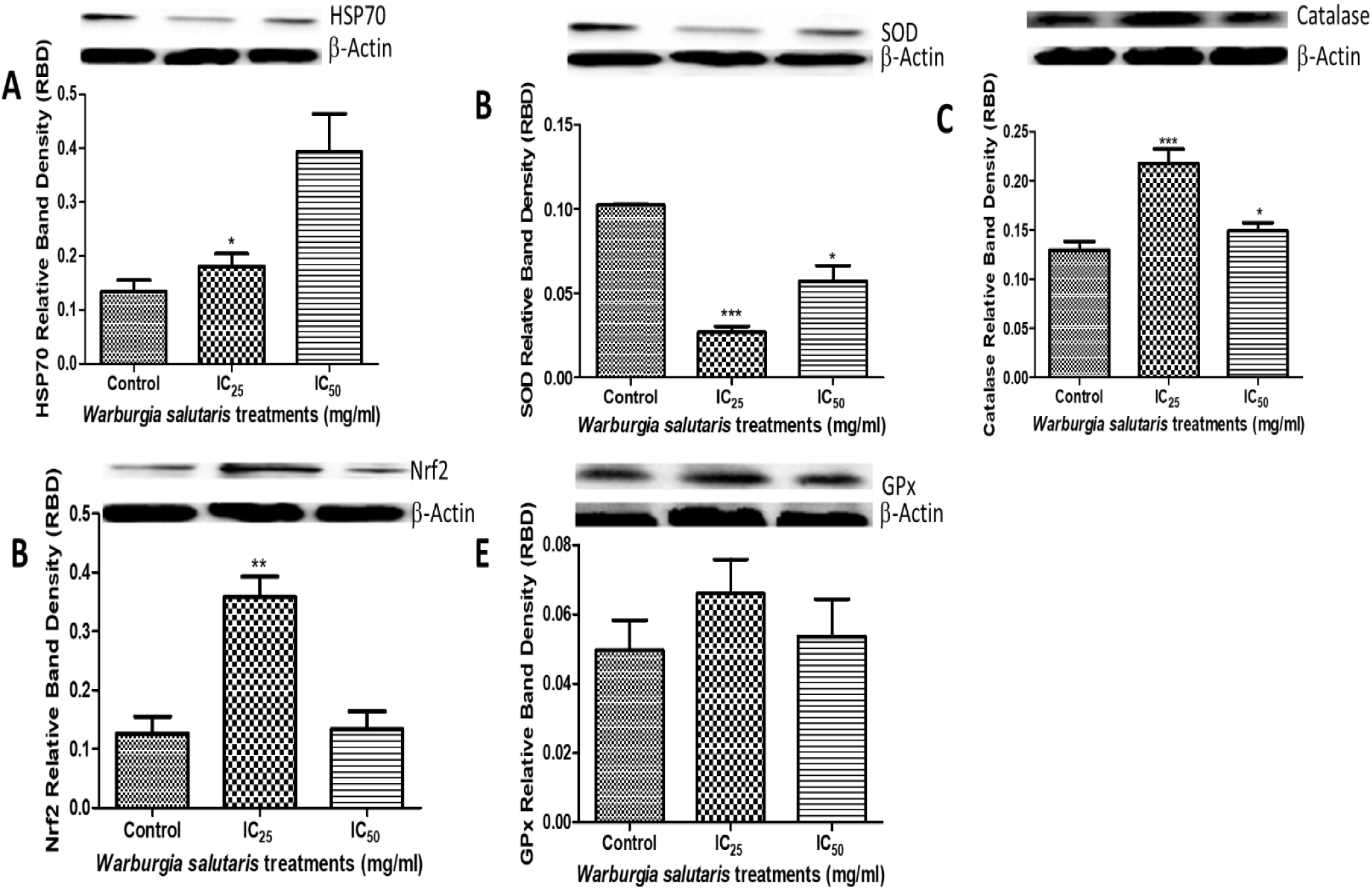
A, B, C, D and E shows oxidative stress-related protein expressions in HepG2 cells treated with *Warburgia salutaris*. A: HSP70 increased significantly at the IC_25_ (\**p*=0.0314) compared to the control. B. Nrf2 protein expression in HepG2 treated cells showing a significant increase at IC_25_ (\*\**p*=0.0026). C. SOD2 protein expression showed a significant decrease at both IC_25_ (\*\*\**p*=0.0008) and IC_50_ (\**p*=0.0141). D. Catalase protein expression was significantly increased at all concentration (IC_25_, \*\*\**p*=0.0005) and (IC_50_, \**p*= 0.0268). E: GPx protein expression in *Warburgia salutaris*-treated HepG2 cells showing a non-significant increase at all concentrations. Students *t*-test with Welch’s correction.

### 3.3 SOD2 gene expression

The SOD2 gene expression (Figure 5a) decreased significantly at all concentrations, with a 2-fold decrease observed for the IC_50_.

**Figure 5a:**
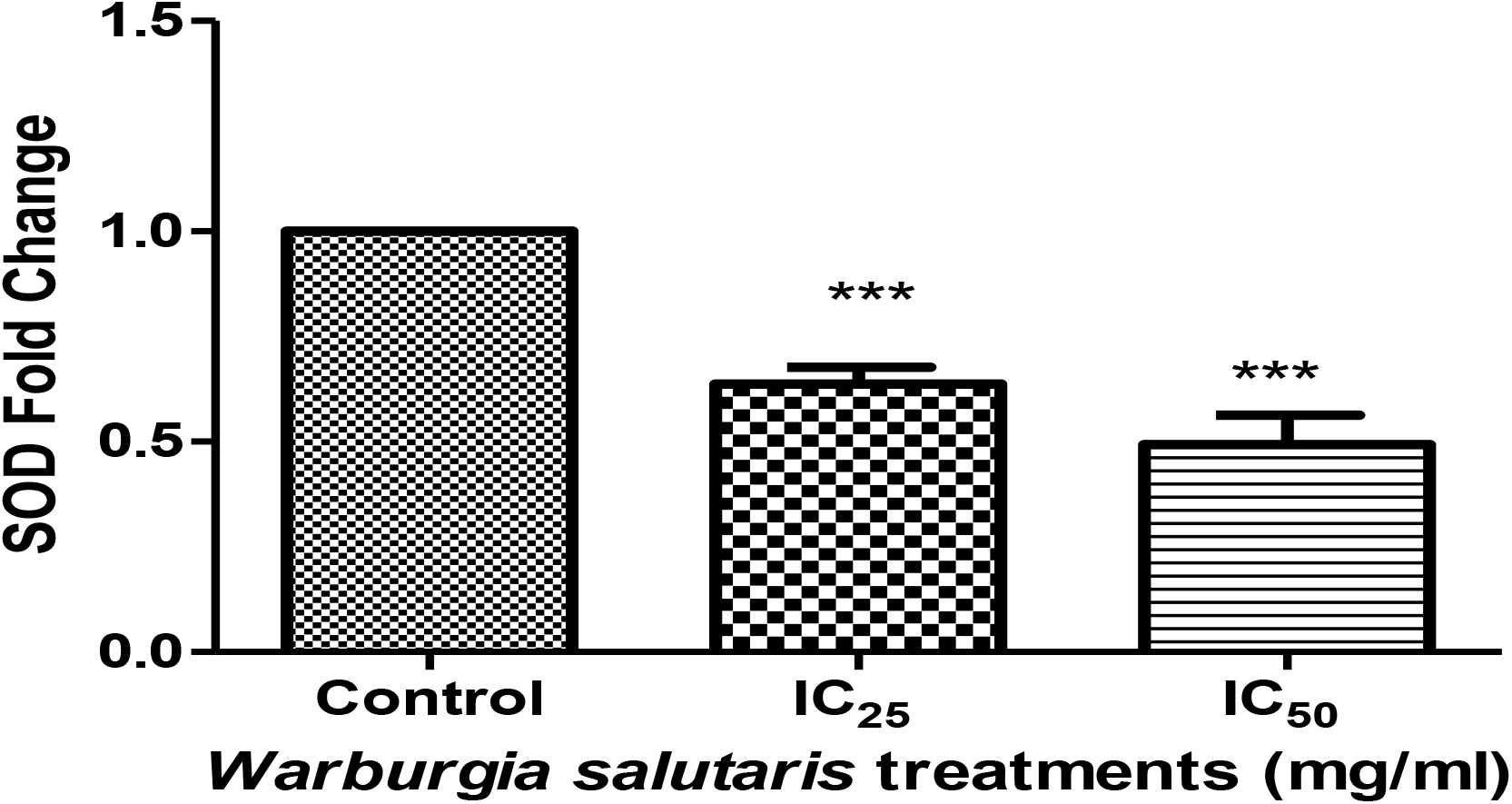
SOD2 gene expression was significantly increased for both *Warburgia salutaris* treatments compared to the control (IC_25_, \*\*\**p*<0.0001 and IC_50_, \*\*\**p*<0.0001); Students *t*-test with Welch’s correction).

### 3.4 Cell death

#### 3.4.1 Apoptosis or Necrosis

Annexin V was used to detect externalised phosphatidylserine, an early marker of apoptosis. Annexin V staining was decreased significantly at the IC_25_, and a non-significant increase was observed at the IC_50_ (Figure 5b).

**Figure 5b:**
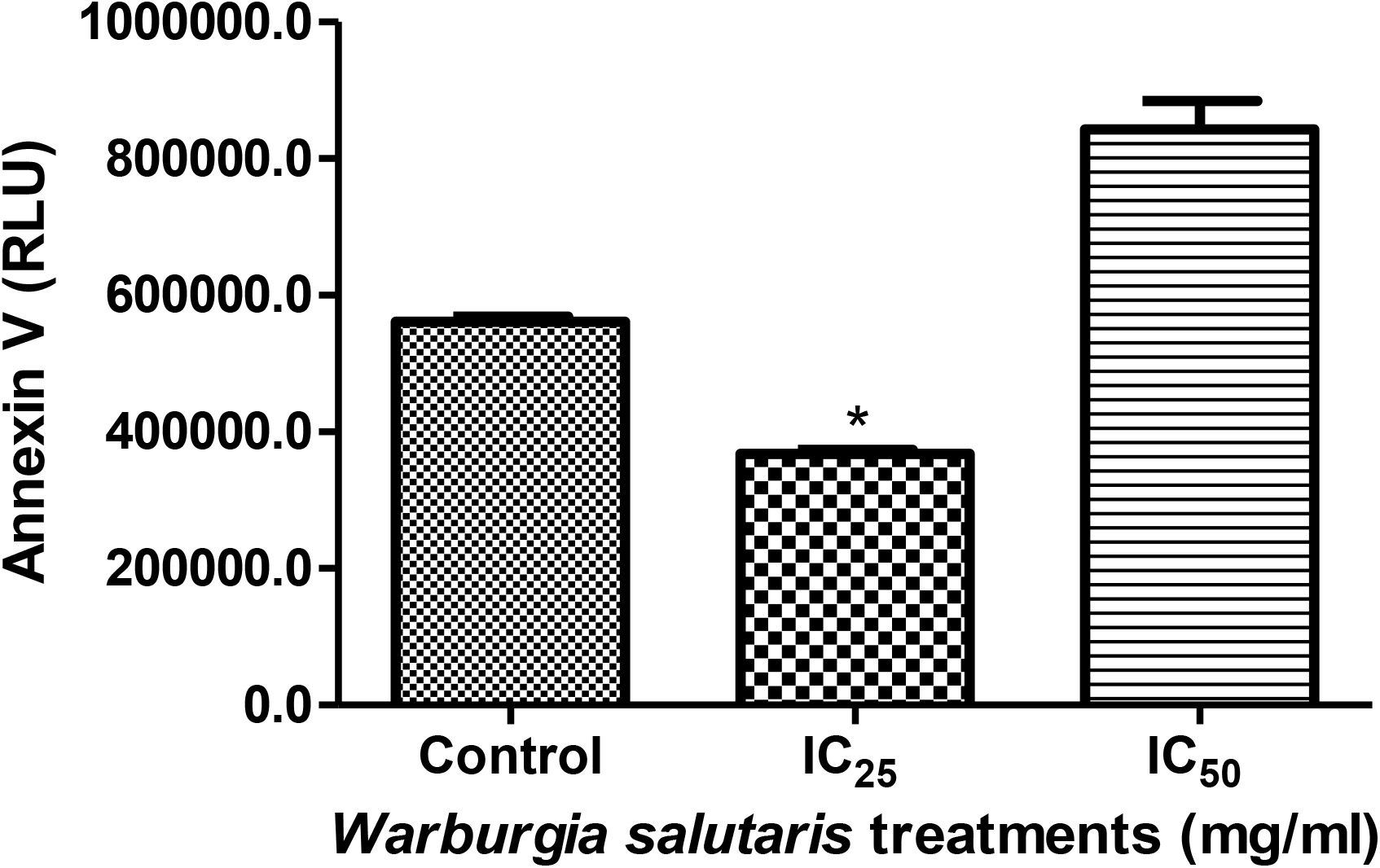
Annexin detection in HepG2 cells treated with *Warburgia salutaris* showed a significant decrease at the IC_25_ (\**p*= 0.0236, Students *t*-test with Welch’s correction).

Cell necrosis was similar to the control at the IC_25_ and a non-significant increase was observed at the IC_50_ when compared to the control (Figure 6A), while LDH decreased significantly (Figure 6B) at all concentrations.

**Figure 6:**
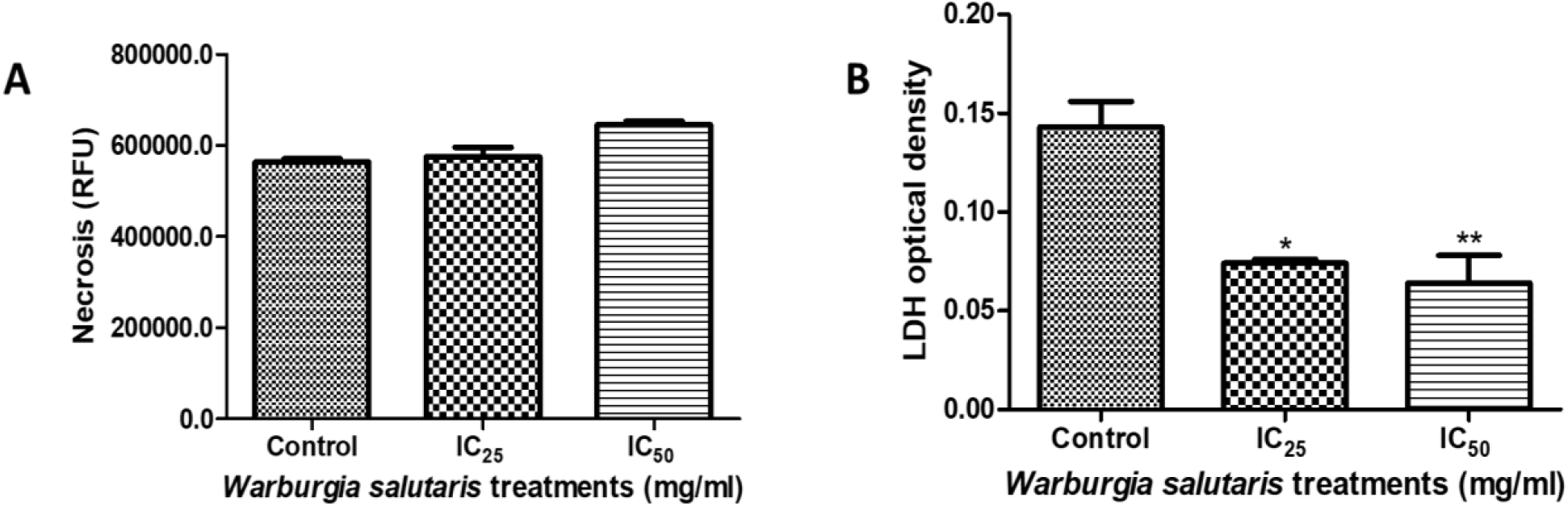
**A.** A non-significant increase in necrosis was noted in *Warburgia salutaris-treated* HepG2 cells. **B** LDH decreased significantly at all concentrations (IC_25_, \**p*=0, 0119 and IC_50_, \*\**p*= 0, 0056; Students *t*-test with Welch’s correction).

#### 3.4.2 Induction and execution of apoptosis: Caspases activity

The induction of apoptosis was assessed by measuring the caspase activities (Figure 7). A slight non-significant increase in caspase 8 activity (Figure 7A) was observed. This contrasted with a non-significant decrease in caspase 9 activity (Figure 7B). Caspase 3/7 (Figure 7C) was increased for both the IC_25_ and IC_50_ treatments, albeit a non-significant change.

**Figure 7:**
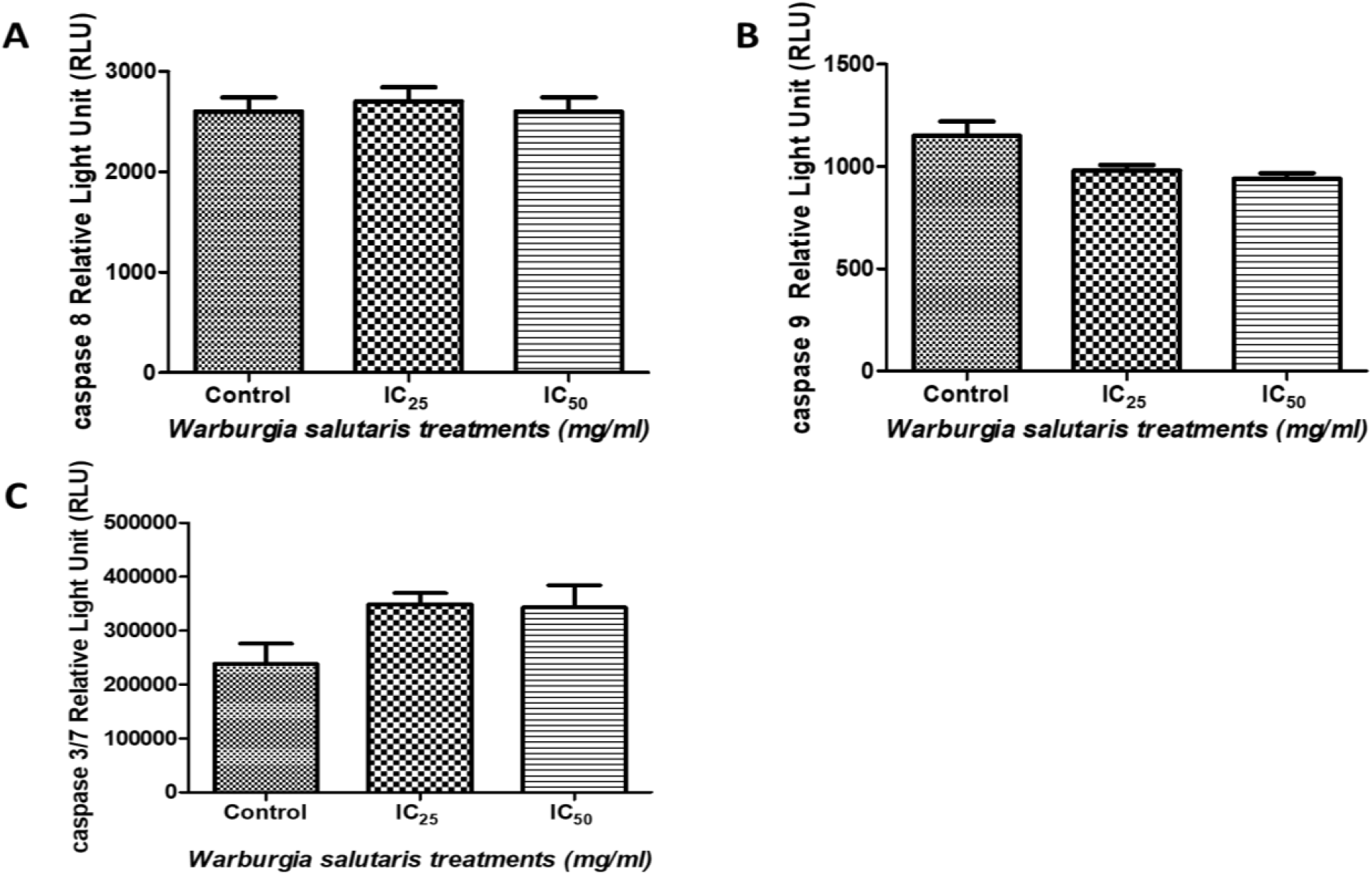
A. Caspase 8 activity in *Warburgia salutaris-treated* HepG2 was similar to the control. B. *Warburgia salutaris* treatment caused a decrease in caspase 9 activity in HepG2 cells. C. Caspase 3/7 activity in HepG2 cells treated with *Warburgia salutaris* was increased relative to the control. All caspases showed no significant difference between the treatment concentrations and the control.

#### 3.4.3. Nuclear morphology

##### 3.4.3.1. Hoechst assay

The Hoechst assay was used to distinguish the different stages of cell division and detect apoptosis. Condensed nuclei, DNA fragmentation, and apoptotic body shape define the late stages of apoptosis. The Hoechst-stained control image (Figure 8A) contains many interphase cells that were stained a uniform blue fluorescence. Brightly fluorescent nuclei depict condensed chromatin in prophase cells, while metaphase (DNA aligned at center ‘equator’ of the nucleus) and telophase (DNA concentrated at the two poles) cells were also visualized (Figure 8A). A decrease in cell density was noted following exposure to *Warburgia salutaris* (Figure 8B, C). The cells containing condensed chromatin were more numerous than in the control.

**Figure 8:**
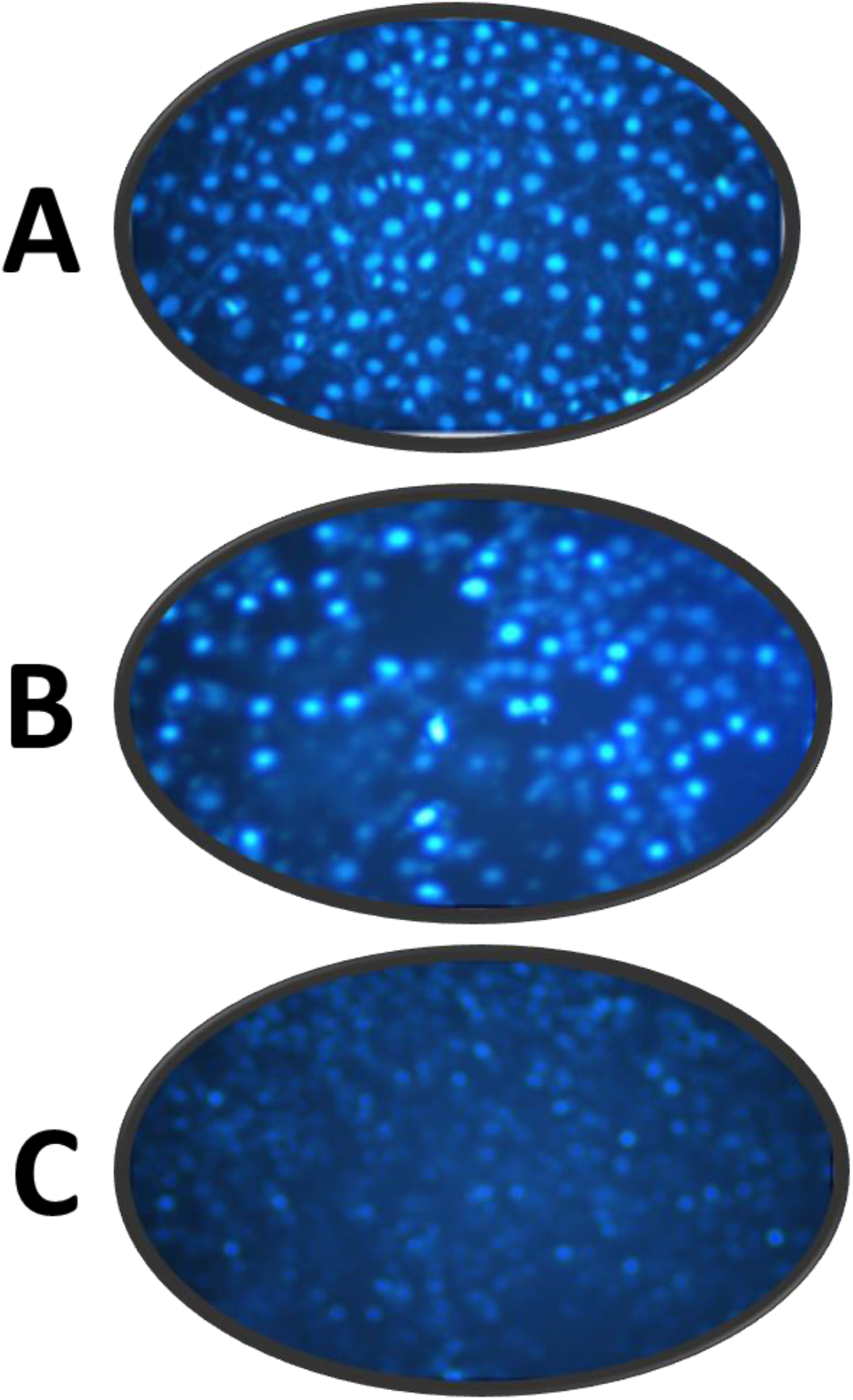
**A.** Untreated HepG2 cells were stained blue, with prophase, metaphase and interphase cells present. **B.** HepG2 cells treated with the IC_25_ concentration of *Warburgia salutaris* show decreased density of interphase cells and numerous cells with condensed chromatin. **C**. The IC_50_-treated cells were sparse with substantial debris present.

##### 3.4.3.2. Comets tail assay

Comets were visualised in all the treatments, including the control. A significant increase in DNA damage occurred in IC_25_-treated HepG2 cells and a notable decrease was observed in the IC_50_ treatment (Figure 9).

**Figure 9:**
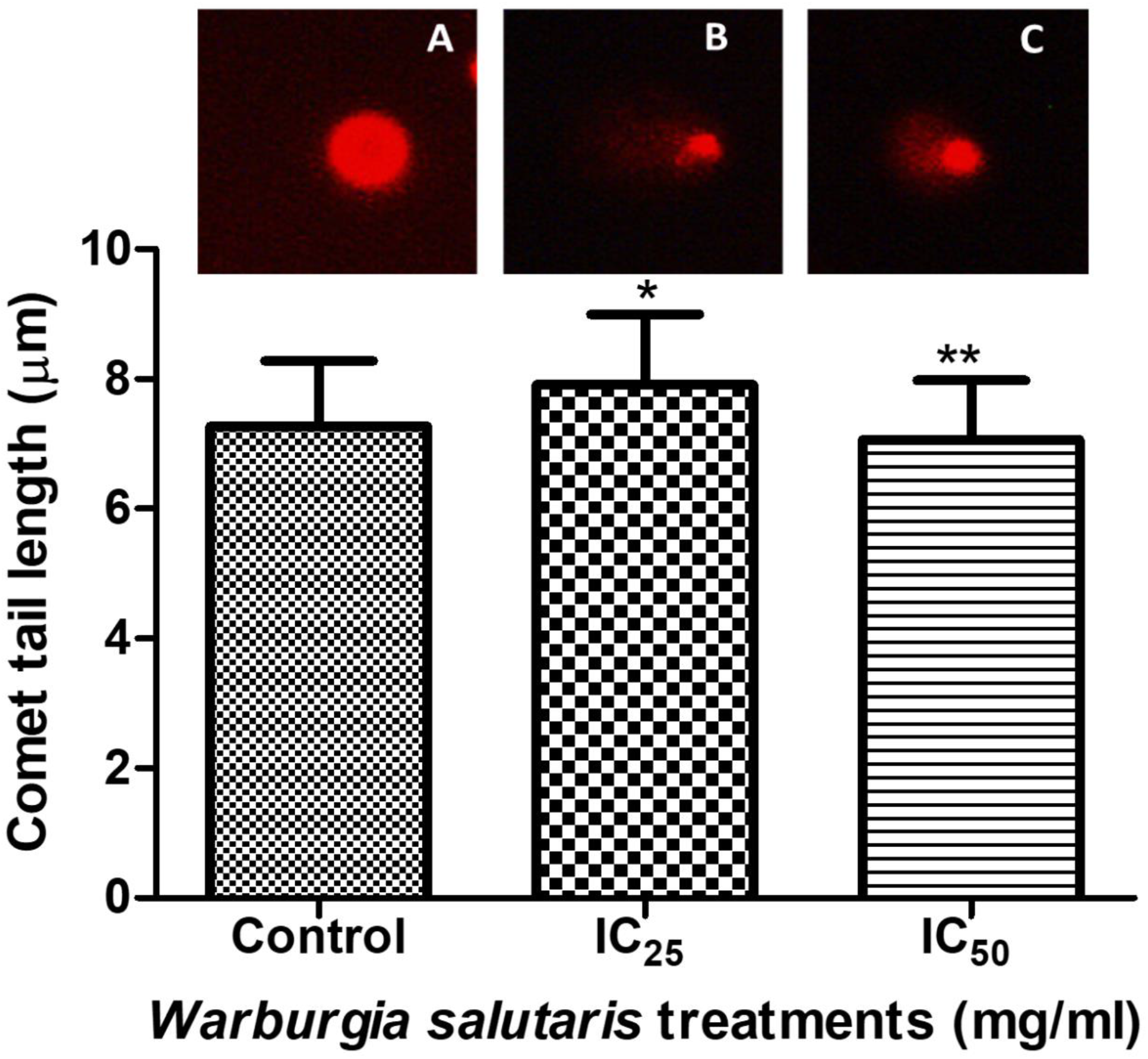
Control cells had small comets (A). Comet tails were slightly increased in length for the IC_25_ treatment (B), and the IC_50_ treatment resulted in a slight reduction in the comet length (C). (IC_25_, \**p*=0.0166 and IC_50_, \*\**p*=0.0062; Students *t*-test with Welch’s correction).

#### 3.4.3. DNA fragmentation

The DNA fragmentation pattern in *Warburgia salutaris*-treated HepG2 cells showed that there was no fragmentation observed at IC_25_ concentrations, while DNA single strand breaks caused fragmentation in the IC_50_ (1.8% agarose gel electrophoresis).

#### 3.4.5. Expression of apoptosis–related protein

Western blotting was used to determine the effects of *Warburgia salutaris* on HepG2 apoptosis-related protein expression (Figure 10). All protein expression was compared to the control. Cleaved PARP protein was significantly increased at the IC_50_; however, the increase at the IC_25_ was non-significant (Figure 10A). Nuclear factor kappa-light-chain-enhancer of activated B cells (NFκB) protein (Figure 10B) significantly decreased at both the IC_25_ and IC_50_, while tumour suppressor protein (p53) increased at the IC_25_ and decreased 11-fold at the IC_50_ (Figure 10C). The apoptosis regulator (Bax) increased significantly at all concentrations (Figure 10D).

**Figure 10:**
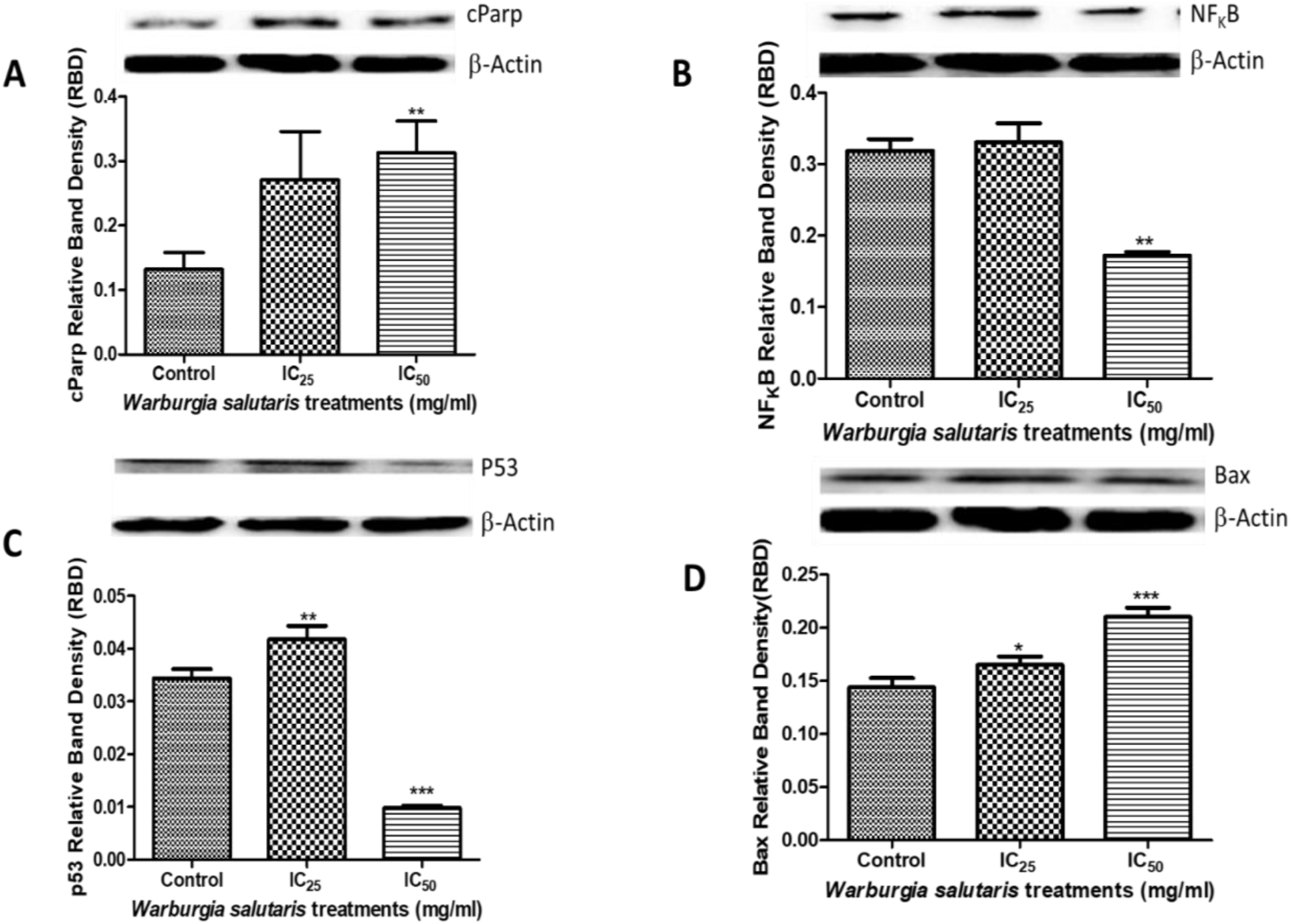
**A, B, C** and **D:** Apoptotic protein expressions in HepG2 cells treated with *Warburgia salutaris*. **A.** cPARP protein expression increased for both treatments, but significantly at the IC_50_ (\*\**p*=0.0030) compared with the control. **B.**NFκB protein significantly decreased at IC_50_ (\*\**p*=0, 0044) **C**: p53 increased significantly at IC_25_ (\*\**p*=0, 0052) and decreased significantly at IC_50_ (\*\**p*=0, 0001). **D**: Bax protein expression increased significantly for both treatments (IC_25_, \**p*=0, 0156 and IC_50_, \*\*\**p*=0, 0001). Students *t*-test with Welch’s correction).

#### 3.4.6. Cell proliferation

Cell proliferation protein markers (Figure 11) showed that the proto-oncogene (cMyc) decreased significantly for all concentrations tested.

**Figure 11:**
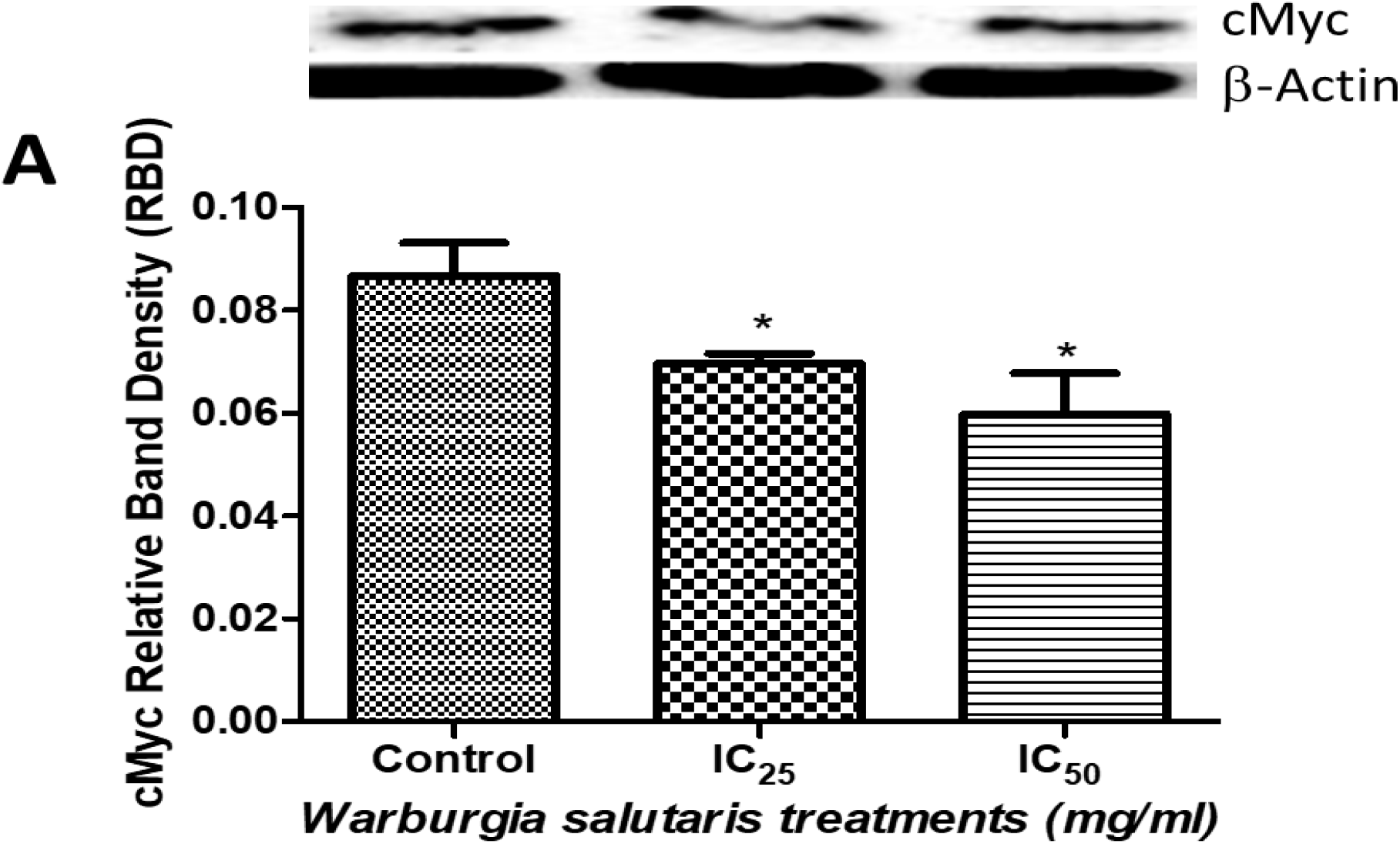
cMyc protein expression showing a significant decrease at the IC_25_ (\**p*=0, 0493) and IC_50_ (\**p*=0, 0197) in HepG2-treated cells compared with the control. Students *t*-test with Welch’s correction.

## 4.0 Discussion

Poor prognosis coupled with high rate of unsuccessful treatment outcome has escalated the burden of liver cancer. This warrant continuous research aimed at the discovery of naturally sourced potently effective anti-cancer treatments. Medicinal plants are popular alternative to cancer chemotherapy as they are inexpensive and easily attainable^17,18^. Native to Southern Africa, *Warburgia salutaris* is among the most valued medicinal plant species globally, used for the treatment of various ailments ranging from common cold and coughs, throat and mouth sores to malaria, venereal diseases, and cancer ^15,19g–21^. To date, the anti-cancer effects of *Warburgia salutaris* has not been fully investigated, and its potential as an anti-proliferative therapeutic in liver cancer has not been explored. Therefore, this study determined the cytotoxic effects of *Warburgia salutaris* aqueous extract in HepG2 cells following an acute exposure (24h).

In this study, the MTT assay yielded an IC_50_ at 4.8mg/ml; similar to the lowest concentration used in a previous study. Furthermore, a dose-dependent decrease in metabolic activity and cell viability of treated HepG2 cells was observed (Figure 1). This finding demonstrates the cytotoxic potential of *Warburgia salutaris* in HepG2 cells and translates to a reduced capacity of the treated cells to convert the MTT salts to a formazan product. This conversion is highly dependent on mitochondrial succinate dehydrogenase and reducing equivalents such as co-enzyme NADH ^22^. The energy transduced in cells is ATP and is produced in viable cells by healthy mitochondria with an intact mitochondrial membrane to ensure the proper functioning of electron transport chain ^23^. Succinate dehydrogenase is embedded in the mitochondrial membrane and forms part of the electron transport chain as well as the Krebs cycle ^24^. It is thus an integral component for ATP generation and indicates a relationship with cellular metabolic activity. The reduced cell viability (Figure 1) in this study correlates with the decrease in ATP (Figure 2) occurring with increased concentrations of *Warburgia salutaris.* This finding is in agreement with a reduction in mitochondrial ATP production attributed to the presence of polygodial, an active compound found in *Warburgia salutaris* ^13^.

The mitochondria are a major source of ROS; they generate O_2_^-^ as a by-product of oxidative metabolism^25^. Under normal physiological conditions, the mitochondria and the cell are equipped with antioxidants that limit the potential damage that can be inflicted by ROS. In the absence of such protection, cells are rendered highly susceptible to the effects of ROS, ultimately resulting in oxidative stress. The MnSOD is responsible for the dismutation of O_2_^-^ to H_2_O_2_. However, in this study both MnSOD protein (Figure 4B) and gene (Figure 5a) expression was reduced following treatment with the *Warburgia salutaris* aqueous extract. As a result, O_2_^-^ cannot be adequately converted to H_2_O, leading to the accumulation of O_2_^-^. Since *Warburgia ugandensis* extracts demonstrate an ability to increase NO levels ^26^, it is probable that the *Warburgia salutaris* extract possesses the same ability. Thus, the simultaneous increased mitochondrial NO production makes NO available to react with O_2_^-^ to yield ONOO^-^, a potent RNS ^27^. In this study, cells treated with *Warburgia salutaris* aqueous extract demonstrated an increase in RNS (Figure 3a), which could potentially nitrosylate lipids and intracellular proteins ^28^.

Results from this study indicated that *Warburgia salutaris* aqueous extract increased MDA concentrations (Figure 3b), suggesting that the extract induced lipid peroxidation and therefore oxidative stress was present within HepG2 cells ^29^. This result concurs with a study carried out by Leshwedi (2008) linking *Warburgia salutaris* methanolic extract and ROS production in mononuclear cells ^13^. It is feasible to suggest that the ingredients of the respective extracts responsible for this effect were present in both the aqueous and solvent extracts. While *Warburgia salutaris* extracts have been shown to suppress lipid peroxidation possibly by scavenging H_2_O_2_ and OH• radicals ^13,30^, the higher concentration used in this study may have induced a pro-oxidant effect.

A concomitant decrease in intracellular GSH (Figure 3d), and an increase in HSP70 (Figure 4A) and Nrf2 (Figure 4D) support the presence of oxidative stress. The decrease in GSH validates the data in Figure 3a, as RNS are known to deplete GSH ^31^. The GSH is critical in the cellular defense against free radicals and hydroperoxides. Its depletion may facilitate the lipid peroxidation observed in Figure 3b that can therefore proceed unhindered. The decreased GSH also indicates that the cells protective capacity may be overwhelmed by the extent of oxidative-related molecular manifestations within the cell. The Nrf2 response may be attributed to the decreased GSH and possibly *Warburgia salutaris* exposure. It could represent the first step by *Warburgia salutaris* polyphenols in the antioxidant response by indirectly upregulating antioxidant expression; Nrf2 protein is a modulator of several antioxidant genes ^32^. However, in this study the increase in Nrf2 protein expression did not induce an increase in MnSOD gene expression (Figure 4D and 4.8). In contrast; both GPx (Figure 4E) and catalase (Figure 4C) that function to detoxify H_2_O_2_ to H_2_O were increased ^33^. Collectively, these results suggest an increase in free radical production and an antioxidant response mounted against it that was inadequate in the prevention of acute oxidative stress.

ROS availability negatively modulates NFκB signaling ^13,34^. The downregulation of NFκB (Figure 10B) correlates with the observed GSH depletion, as shown in human leukaemia and prostate cells ^35^. Meng *et al.* (2010) suggests that this relationship may be exploited to improve the effectiveness of therapeutic strategies against cancer^35^. Mononuclear cells also slightly downregulated NFκB in response to *Warburgia salutaris* exposure, possibly by exploiting its antioxidant potential and blocking delivery of oxidants to the cell or modification of the intracellular redox state^13^. The latter is more likely in this study in light of the increased free radicals. The interference with redox homeostasis potentially disrupts cell survival pathways, thus steering the *Warburgia salutaris* treated cells toward cell death ^36^.

Cell death via apoptosis may proceed using the intrinsic, extrinsic or endoplasmic reticulum pathways ^37^. Externalisation of phosphatidylserine (Figure 5b) provides evidence that apoptosis occurred at the IC_50_. Additionally, PI fluorescence was slightly increased (Figure 6A) but was accompanied by diminished levels of LDH (Figure 6B) suggesting that necrosis was unlikely the cause of cell death. In fact, combined PS and PI results may indicate the presence of late stage apoptosis ^38^. The cell death that occurred in *Warburgia salutaris* treated cells (Figure 8B, C) may therefore be attributed to apoptosis. To maintain homeostasis, cell death processes are often counterbalanced with cell proliferation. However, reciprocal cell proliferation was not evident as shown by a decrease in cMyc (Figure 11).

The workers of apoptosis are caspases. Caspase 8 (Figure 7A) was not affected by *Warburgia salutaris,* thus apoptosis was not via the extrinsic pathway. An observed increase in Bax protein expression (Figure 10D) suggests that caspase 9 should be activated; however, decreased caspase 9 activity (Figure 7B) in this study correlates with increased HSP70 (Figure 4A); alternatively, nitrosylation of caspase 9 by RNS (Figure 3a) may also contribute to the reduced activity ^28^. Therefore, the increase in caspase 3/7 (Figure 7C) that confirms that cells were primed for execution of apoptosis may be attributed to alternate mechanisms for caspase 3 activation, including granzymes and/or the endoplasmic reticulum pathway ^39^.

The main target for caspase 3/7 is Parp-1, which is cleave into two fragments (cPARP). cPARP (Figure 10A) was increased after exposure to *Warburgia salutaris*. Another hallmark of apoptosis facilitated by caspase3/7 is DNA fragmentation that is accomplished by caspase-activated DNase (CAD) ^38^. Comet tails were not markedly increased (Figure 9), indicating minimal DNA fragmentation. Leshwedi (2008) also showed decreased comets in response to 5mg/ml *Warburgia salutaris* ^13^. The absence of DNA aberrations is reinforced by the decrease in p53 (Figure 10C) which detects DNA damage and activates DNA repair mechanisms, and decreased OGG1 (Figure 3c), a repair enzyme for oxidative DNA damage. There is evidence that *Warburgia salutaris* extracts contain mannitol, a potent OH• scavenger, which together with flavonols and flavonoids may be responsible for reduced DNA damage in *Warburgia salutaris* exposed cells.

## 5.0. Conclusion

The study investigated the molecular mechanisms associated with the effects of *Warburgia salutaris* on human hepatoma cells. The findings show that *Warburgia salutaris* modulates oxidative stress and apoptosis in HepG2 cells. The disruption of redox homeostasis and activation of apoptosis may indicate the potential of *Warburgia salutaris* as an anti-cancer agent that would serve as an alternative to conventional therapeutic agents. This is particularly important for the protection of these valuable medicinal plants in their natural environments, as it is among the International Union for Conservation of Nature (IUCN) Red List of endangered species. However, further studies on *in-vivo* animal models may be done to verify the results.

## Ethical approval

Ethical approval was obtained from the Biomedical Research Ethics Administration under the Reference number: **BE036/19**.

## Acknowledgements

The author is grateful for financial assistance from the National Research Foundation and College of Health Sciences.

## Data Availability Statement

The generated data used to support the findings of this study are included within the article.

